# Making data-driven hypotheses for gene functions by integrating dependency, expression, and literature data

**DOI:** 10.1101/2020.07.17.208751

**Authors:** Matthew D. Hirschey

## Abstract

Identifying the key functions of human genes is a major biomedical research goal. While some genes are well-studied, most human genes we know little about. New tools in data science -- a combination of computer programming, math & statistics, and topical expertise -- combined with the rapid adoption of open science and data sharing allow scientists to access publicly available datasets and interrogate these data *before* performing any experiments. We present here a new research tool called data-driven hypothesis (DDH) for predicting pathways and functions for thousands of genes across the human genome. Importantly, this method integrates gene essentiality, gene expression, and literature mining to identify candidate molecular functions or pathways of known and unknown genes. Beyond single gene queries, DDH can uniquely handle queries of defined gene ontology pathways or custom gene lists containing multiple genes. The DDH project holds tremendous promise to generate hypotheses, data, and knowledge in order to provide a deep understanding of the dynamic properties of mammalian genes. We present this tool via an intuitive online interface, which will provide the scientific community a platform to query and prioritize experimental hypotheses to test in the lab.

## Introduction

Identifying the mechanisms by which genes interact to control biologic processes is a major biomedical research goal (Barabasi et al., 2011). Historically, most research studies interrogating the function of a gene have taken a one-gene-at-a-time approach. More recently, combinatorial gene deletion studies have revealed synergistic or epistatic functional gene-gene associations, such as in yeast (Tong et al., 2004) and in human cells (Horlbeck et al., 2018).

However, moving from large-scale genetic maps to candidate annotations for the molecular functions of individual genes remains a challenge. Predictive analytic approaches via computational biology and data science are increasingly used to overcome this challenge. As one example, early studies in yeast showed that proteins functioning within the same process are more likely to result in similar phenotypes when their genes are disrupted (Fraser and Plotkin, 2007). In mammalian cells, the development and use of sophisticated experimental and data processing pipelines have established genome-wide functional interaction networks, which can be interrogated as publicly available datasets (Kim et al., 2019; Meyers et al., 2017). Thus, predictive analytical approaches to map gene function have the potential shed light on hundreds of human genes and associated functions.

One simple example of this approach is using gene co-expression analysis. When a gene of unknown function is identified, one strategy to learn something about the new gene is to identify shared patterns of expression with other genes. If unknown Gene X is expressed with known genes A, B, and C, then you can infer that Gene X might be part of a functional module with A, B, C (Figure 1a). This approach is particularly powerful when genes A, B, and C are co-expressed as part of a known biological pathway, which leads to the hypothesis that Gene X might also be part of that same pathway. This approach leverages the heterogeneity present within a broad range of cells and that coordinated behavior of gene expression is consistent across cells.

Correlations of patterns of gene expression can be extended to other cellular behaviors, such as gene essentiality (Figure 1b). The power of this idea – sometimes called co-essentiality mapping – lies in using a functional read-out to map genes to common functional pathways as opposed to using gene expression as a read-out (Wang et al., 2017). In the updated example (Figure 1b), genes that are in the same functional pathway will result in similar patterns of dependency across cell lines. In this example, if one set of cell lines requires a pathway for survival (it is a pathway containing essential genes), but not another set of cell lines, then identifying genes that share similar patterns of dependency can be revealed (Figure 1b). These patterns of similarity can then be prioritized for experimental interrogation. Importantly, this approach has been successfully used to identify novel gene functions (Bayraktar et al., 2020; Pan et al., 2018; Wang et al., 2017)

Project Achilles is a systematic effort by the Broad Institute as part of a larger ‘DepMap’ project aimed at identifying and cataloging gene essentiality across hundreds of well-characterized cancer cell lines using standardized, pooled, genome-scale loss-of-function screens. This project uses lentiviral-based pooled CRISPR/Cas9 libraries to systematically knock-out each gene in the genome, which allows for the stable ablation of each gene individually in a subset of cells within a pooled format allowing for genome wide interrogation of gene essentiality. Using computational modeling, a normalized value of gene essentiality is given for each gene in a single cell line. A lower score means that a gene is more likely to be essential in a given cell line. A score of -1 corresponds to the median of all common essential genes, whereas a score of 0 is equivalent to a gene that is not essential; a positive score indicates a gain in fitness and often identifies tumor suppressor genes. The ‘DepMap’ provides a rich source of information that can reveal similar genetic dependency patterns between genes, implying functional relationships.

We envisioned that when integrated with other information, hypotheses about relationships between identify novel and unexpected genes can be generated. Thus, we set out to map genes to common functional pathways based on dependence of a pathway for cell viability.

## Results

### Predictive analytics by pattern recognition

First, essential gene data from Project Achilles were downloaded from the DepMap portal at: depmap.org. The 20Q1 release contains gene essentiality scores for 17426 genes across 739 cell lines (Figure 2a). To find patterns in gene dependencies across cell lines, we generated a Pearson correlation matrix of all genes by all genes, as has been done previously (Barido-Sottani et al., 2019). This analysis generated gene-gene correlation values that matched values published on depmap.org, validating the first step in our analysis.

High levels of gene expression are often thought to be indicative of key genes for a given cell type. Thus, we next compared dependency values to gene expression values. The Cancer Cell Line Encyclopedia project is a collaboration between the Broad Institute, and the Novartis Institutes for Biomedical Research and its Genomics Institute of the Novartis Research Foundation, which together conduct detailed genetic and pharmacologic characterization of a large panel of human cancer models. As of the CCLE 2019 release, 1270 cell lines have been characterized for gene expression. In the 739 cell lines from the 20Q1 DepMap release, 733 have gene expression data. Using these two datasets, we compared the essentiality of a gene to its expression value (Figure 2b). We predicted a V-shaped curve, with stronger dependencies as gene expression increases. Surprisingly, we saw no relationship between gene expression and gene essentiality, where genes with both low and high expression displayed both gains and losses in fitness. The overall observation from this dataset shows baseline gene expression levels are poor indicators of the essentiality of a gene.

This analysis also highlighted that several genes were binned on the x-axis, i.e. could have no measurable expression levels, but have assigned dependency scores. Across 739 cell lines in the Achilles project, 16.8% of all gene expression values are zero (Figure 2c), confirming this notion.

### Noise Reduction

Our goal was to construct functional gene networks to learn more about cellular processes. We reasoned that genes not expressed in cells would be unable to participate in these functional networks. Given cells do not express all genes, but might receive a dependency score in the experimental paradigm, we sought to remove dependency scores for gene-cell line pairs that have an expression value of zero under basal conditions. From the 733 cell lines with both expression and essentiality data, we removed dependency scores for genes from cell line that have a corresponding gene expression value of zero.

For some genes expressed in highly specific and restricted cell types, this operation removed many dependency values. After removing these values, we found that highly specialized genes in discrete cell types have too few cells with both gene expression values and gene essentiality values to assign a meaningful correlation value. Thus, if a gene was absent from too many cell lines, we omitted it to prevent assigned values from relying on too few data points (Figure 2d).

We set a threshold to remove the bottom 5% of the genes with the fewest number of cells deriving the correlation value and having the highest number of correlated gene patterns (Figure 2e). This threshold removed genes that were present in fewer than 108 cell line and missing in more than 631 cell line. We reasoned that the correlation pattern of a gene would be meaningless if the data was derived from too few cell lines, and that gene was therefore removed. This process removed 908 genes that had too few cells with expression and dependency data.

To quantify the effect of these cleaning steps on the data, we compared the correlation matrix generated from the unprocessed data present in the DepMap to the correlation matrix from the cleaned data here. Plotting these ∼200M original-cleaned gene pairs revealed that more positive genes became more positive compared positive genes that became negative; conversely, more gene pairs with negative correlations became more negative rather than less negative (Figure 2f). Simply stated, these cleaning steps had a greater effect on strengthening the data in either direction rather than weakening it, supporting these steps as important for this resource. These ‘cleaned’ dependency data were then used to generate correlation matrix.

To identify genes that shared similar patterns of essentiality with other genes, thereby placing genes in functional pathways, we generated a gene-by-gene Pearson correlation matrix on these prioritized data to quantify the similarity in dependency patterns and annotate genes in functional pathways. This process generated approximately 304 million correlation values, with a distribution centered around zero (Figure 3a) and produced a range of maximum correlation values for each gene (Figure 3b).

### Thresholds

Rather than setting an arbitrary threshold for the r^2^ value that would be considered a low, medium, or high correlation between two genes, we performed a permutation test on the correlated data. We sampled 20,000 r^2^ values from all gene-gene pairs without replacement simulating a virtual Achilles correlation dataset for a single cell line. We then repeated this process 1000 times mimicking 1000 discrete cell lines. This analysis produced a mean correlation of 0.003 and a standard deviation of 0.069.

Using a standard deviation threshold of 3, we calculated the boundaries of r^2^ values to be greater than 0.21 or lower than -0.20 for negative correlations. Simply stated, gene-gene correlations above or below, respectively, these values represented ∼1% of the dataset, were considered strong, and therefore used for subsequent analyses.

### Comparing to Known Literature

The process of generating a prioritized gene list based on similar patterns to a query gene is an unbiased approach to map genes into functional pathways. To find novel genes in established pathways, this approach requires identifying “novel” or “unexpected” genes in the gene list. However, what may seem unexpected to one researcher might be obvious or intuitive to another. To remove bias in prioritizing candidate genes for further study and to programmatically identify novelty, we generated a co-publication index, where every human gene was compared to publication co-incidence with every other human gene in the PubMed database.

A new machine learning (ML)-based resource developed by the Computational Biology Branch at the NLM/NCBI called Pubtator aims to overcome challenges in gene nomenclature by using advanced text-mining techniques to develop a ML model that generates automated annotations for genes/proteins, genetic variants, diseases, chemicals, species and cell lines within a single paper, across all PubMed (30 million) abstracts and a PMC Text Mining subset of (3 million full text) articles (Wei et al., 2019). Using these data, we performed a pairwise count of each gene with each other gene in a single paper, across all papers, thereby determining the number of times each gene had been co-published with another gene. Because of the wide range of annotated gene co-occurrences (0 to 23197), we also generated a relative index normalized to 100 [(co-occurrence count/max co-occurrence)*100]. The co-occurrence count and co-occurrence index provide useful, and unbiased information about the level of prior knowledge for gene-gene relationship identified by the analytic approach detailed above.

### Pathway Analyses

To identify clusters of co-essential genes with shared relationships, we performed gene set enrichment analysis. Enrichment analysis is a computational method for inferring knowledge about a target gene set by comparing it to annotated gene sets representing prior biological knowledge. For each gene in our matrix, we determined the number of genes that were greater than or less than 3 standard deviations away from the permuted mean correlation. This target gene list was then queried against 108 gene sets across a broad range of curated data. By leveraging the Enrichr resource (Kuleshov et al., 2016), we determined the top ranked pathways, processes, drugs, cell lines, tissues, or diseases, and ranked by p-value.

### Network Analyses

As an alternative approach to prioritize further study of novel functional mappings, an interactive network graph of gene-gene relationships was built. A single gene query results in a list of top and bottom associated genes based on patterns of gene dependency scores. The top 10 and bottom 10 genes are then queried for each of their top and bottom correlated genes.

These resulting 400 gene pairs (20 genes * 20 genes) are then used to build a network graph, which can reveal if a top associated gene with the query gene, also has the query gene in its top (or bottom) 10. Genes (nodes) with only one relationship (edge) were pruned to simplify the network.

### Data Access and User Interface

We present these analyses as a comprehensive research platform for predicting pathways and functions for thousands of genes across the human genome. We provide an intuitive online interface located at www.datadrivenhypothesis.org, which allows users to query and prioritize experimental hypotheses to test in the lab. In order to identify the functional annotation of a single gene, begin with a query.

Entering a single gene in the search box produces a series of tables and plots that identifies a functional map of the processes that gene might be involved in. In some cases, querying a gene with known functions will identify genes with well-established connections to the query gene; in other cases, new genes and new biological process might be identified, suggesting there is more to discover for well-known pathways. Querying unknown genes is especially powerful, as the associated genes and pathways provide a starting point for an otherwise difficult problem to prioritize experimentally.

### Query Example

As an example, we queried the protein P53. Querying *TP53* (official gene symbol) generates a short summary of the gene, its name and list of aliases, and an Entrez gene summary paragraph when available.

The first plot shows the distribution of dependency scores across 739 cell lines ranked from lowest (strongest dependencies) to highest (no dependency or inverse) (Figure 4a). Each of the 739 cell lines is represented by a single point on the plot. Generally, values below -1 indicate the gene of interest (*TP53* in this example) is essential in that cell line; values between - 1 and 0, mean that cells lose fitness, but the gene is not essential; values hovering around zero indicate that ablation of *TP53* has little effect on cell growth; values above 1, indicate that knocking-out the gene leads to a fitness advantage. In the case of *TP53*, several cells have a fitness advantage in its absence, consistent with its role as a tumor suppressor. The online version of this plot is interactive, allowing users to quantify the histograms. The second plot is a histogram of dependency scores, showing the distribution of scores for *TP53* (Figure 4b). While the majority of cells have little change in cellular growth when *TP53* is absent (the histogram is centered around zero), some cells require TP53 for growth (cells scoring below -1), whereas in other cells *TP53* functions as a tumor suppressor (cells with a score above 1). The online version of this plot is interactive, allowing users to zoom, query, and identify specific cell lines.

The third and fourth plots are horizontal boxplots of cell lines grouped by lineage or sub-lineage.

To identify the cells at the tails of the plots, a dependency table shows the ranked cells by dependency score with cell lineage information appended. In some cases, specific cell types or lineages will show consistent patterns of dependency on a gene. Understanding the shape of the curve and distribution of the raw data underlying the patterns is important for interpreting the results.

Positive correlations of dependency scores are ranked for each gene, which show similar patterns of dependencies in the same cell lines. More simply, the cells that care about *TP53* deletion also care about deletion of these genes, implying a functional relationship. In the Dependency Score Example heatmap schematic above, *TP53* is gene X, and genes with similar patterns would be genes A, B, and C. The 129 genes that show a similar genetic dependencies as *TP53* and are above 3 standard deviations away from the resampled mean are shown. One data source that can help prioritize genes for study is to compare gene-gene associations to known gene-gene relationships based on previous literature. By counting the number of times p53 was published with each of these top ranked genes, the Relaxin family peptide receptor 2 (*RXFP2*) emerges as an unexpected connection, based on current literature, and could be prioritized for further interrogation. These 129 genes were queried for gene set enrichment, and the gene sets and pathways with the strongest statistical significance are shown. Simply stated, these are the pathways that best represent the list of genes that share similar genetic dependencies, and suggest that the query gene is part of these pathways.

Like the analysis for genes that share similar patterns, this analysis can be used to find genes that share distinctly dissimilar patterns; that is, genes that have an inverse correlation of dependences. Simply stated, the cells that care about *TP53* deletion do not care about deletion of these genes, implying an inverse or opposing relationship. In the Dependency Score Example heatmap schematic above, *TP53* is gene X, and genes with dissimilar patterns would be genes D, E, and F. The 186 genes that show inverse genetic dependencies to *TP53* and are below 3 standard deviations away from the resampled mean are shown. These 186 genes were also queried for gene set enrichment, and the gene sets and pathways with the strongest statistical significance are shown. Simply stated, these are the pathways that best represent the list of genes that have inverse genetic dependencies. The interpretation of these genes and pathways is more variable than the positively correlated genes and pathways. In some cases, a negative regulator of a gene has a negative correlation with that gene, such as in this example with *TP53* and its negative regulator *MDM2*. In other cases, opposing pathways are shown, contrasting the *TP53* enriched pathway term “RB Tumor Suppressor/Checkpoint Signaling in response to DNA damage Homo sapiens h rbPathway” with the dissimilar enriched pathway term “ribosome biogenesis (GO:0042254)”, revealing two opposing biological pathways.

Identifying genes that share similar patterns of dependency to a queried unknown gene generates strong hypotheses about new functional annotations and maps to new pathways. However, the strength of the hypothesis cannot be fully inferred from single gene list. If a new gene is associated with the queried gene, then you might infer a functional relationship.

However, if you inspect the top 10 genes with the queried gene, then inspect the top 10 genes of each of those, building a functional network graph of the top related genes might reveal a stronger association of the new gene with your queried gene and its top ranked genes (Figure 4c). The data presented on datadrivenhypothesis.org is an interactive image that can be zoomed, dragged, and manipulated for data exploration.

## Discussion

It is well-known that human cancer cell lines rely on different pathways for their viability. Indeed this is the entire rationale for personalized, precision medicine in cancer. The overall goal of the ‘DepMap’ project is to identify all essential genes in 2000 cell lines over the 5-year project period to identify new therapeutic targets in various cancers. Despite not knowing the mechanistic basis for why some cell lines require specific genes while other cell lines do not, we reasoned that intrinsic reliance of a cell on a pathway might allow unbiased detection of novel genes participating in specific pathways.

DDH represents a comprehensive resource for predicting pathways and functions for thousands of genes across the human genome. Importantly, this method integrates gene essentiality, gene expression, and literature mining to identify candidate functions of known or unknown genes. We present this resource via an intuitive online interface, which provides the scientific community a platform to query and prioritize experimental hypotheses to test in the lab.

We started with several high-quality, functional genomic datasets. Functional genomics is a field of molecular biology that aims to understand the function of all genes and proteins in a genome – a stated goal of much basic science research. In functional genomics, experimental strategies generally involve high-throughput, genome-wide approaches rather than a more traditional “gene-by-gene” approach. The advent and rapid adoption of data-sharing platforms made available with Creative Commons Attribution 4.0 International (CC BY 4.0) licenses have provided high-quality data sets for public interrogation.

Mapping genetic interactions from these datasets can be performed using a variety of different data types. Gene co-expression values, methylation patterns, ChIP Seq, and ATAC Seq data have all been used to assign interactions based on gene expression. Similarly, broadscale functional genomic screens, such as with CRISPR-Cas9, have largely relied on cellular fitness (growth and proliferation) as the basis on which to build functional networks. Here, we have used sgRNA depletion-based and copy number corrected gene essentiality score data from Project Achilles to define genetic networks based on patterns of changes in cellular fitness on a spectrum of essential (gene ablation results in cell death) to prohibitive for cellular growth (gene ablation results in growth advantage) (Meyers et al., 2017).

A major consideration when inferring gene functions from parallel genome-wide screens is the processing method used to identify meaningful associations and remove confounding (i.e. ‘noisy’) data. In this analysis, we approached data processing in two steps. First, under the rationale that genes with no expression cannot participate in a biochemical function, we removed all gene essentiality data in a cell line that had an expression level of zero in that same cell line. The second step in our workflow removed genes that were expressed in too few types of cells to be representative of the entire dataset. False positive gene-gene interaction hits such as off-target effects of sgRNAs been reported (Fortin et al., 2019; Kim et al., 2019). Our approach to strengthen the data resulted in more positively and more negatively correlated gene-gene pairs.

While the top correlated genes from dependency score data is available on depmap.org, a unique strength of the DDH pipeline is its ability to build genome-wide interactive networks based on similarities in gene dependency scores, while also considering gene expression and previous knowledge based on data mining. Genome wide CRISPR screens have previously been used to construct protein-protein interaction networks (Pan et al., 2018), and identify synthetic lethal gene interactions (McDonald et al., 2017; Wang et al., 2017).

In DDH, we leveraged high-quality full-genome CRISPR screens of over >700 cancer cell lines representing a diverse range of cancer sub-types across most tissues in the human body. The Achilles resource was originally developed for precision cancer medicine; however, identifying patterns of similar behaviors across these cell lines can reveal fundamental gene-gene interactions beyond cancer. This approach leverages intrinsic heterogeneity between the profiled cell lines that have discrete genetic liabilities. The high number of profiled cells allows mapping specific genes to functions based on similar patterns of liabilities across lines, without asking *why* each cell has a specific liability.

Despite the presence of several ‘positive controls’ in the data (i.e. identifying known functional relationships between genes), we’ve also identified several genes for which this resource provides no functional mapping. For example, if a cell-type-specific gene was expressed in too few cells, then the correlation values were artificially elevated. In the extreme case, some genes were expressed in fewer than 5 cell lines, and were among the strongest signals in the data set – an artifact of having too little data. Furthermore, we found clusters of genes that have no strongly correlated genes, but high median correlation values. The small range of essentiality values driving the correlations in this set of genes weakens the predictive capacity, as has been described previously for olfactory receptors (Boyle et al., 2018).

We find the easiest way within DDH to determine the predictive utility of functional gene mappings is to first build a network graph for the gene of interest. If a strong, connected network emerges, where the target gene is strongly correlated with several of the top connected genes, then a functional module exists. Alternatively, if the target gene is not present on the graph, then it is not connected in a network to its top genes, and DDH has little predictive utility for the target. Second, if the target gene is in a strong network, then inspecting the co-publication count and index could reveal unexpected members in functional gene modules.

To ensure access to this resouce, we developed an intuitive online interface that allows exploration and interrogation of the data underlying DDH. Importantly, we built a development pipeline that allows facile updating corresponding to quarterly releases of the ‘DepMap’ project, as well as yearly data releases of literature-minded data, together ensuring timely and accurate predictions based on the current state of knowledge.

## Conclusion

Mapping a broad set of gene functions *en masse* is currently not possible, and therefore most scientific research proceeds with a one-gene-at-a-time approach. While this reductionist approach has guided the scientific method for hundreds of years, the volume, complexity, and sophistication of modern science necessitate alternative approaches.

Like the proverbial man searching for lost keys under the lamp post because the light shines there, searching for biological truths often occurs under ‘lamp posts’ because that’s where scientists can see. New tools in data science combined with the rapid adoption of open science and data sharing allow scientists to access publicly available datasets and interrogate these data before performing any experiments. Indeed, these analytical tools allow researchers to identify new hypotheses for well-studied genes, or new processes for un-annotated genes.

The scientific method has guided scientific minds for hundreds of years, starting with a question, followed by a hypothesis, and then an experimental path to test the prediction. While hypotheses are the bedrock of science, the volume, complexity, and sophistication of modern science necessitate new methods to generate hypotheses. We envision resources like DDH will facilitate scientists having strong data to support new hypotheses before testing, and have the potential to accelerate biological discovery.

## ACKNOWLEDGEMENTS

We would like to acknowledge Derek K. Zachman and members of the Hirschey laboratory for discussion and comments. We want to thank John Bradley, Dan Leer, and the Center for Genomics and Computational Biology at Duke University for computational and technical support. We would like to acknowledge funding support from the Glenn Foundation, the National Institutes of Health and the NIDDK grant R01DK115568, the National Institutes of Health and the NIA grant R01AG045351. The content is solely the responsibility of the author and does not necessarily represent the official views of the National Institutes of Health or other funding sources.

## METHODS

### Data Download

First, essential gene data from Project Achilles were downloaded from the DepMap portal at: depmap.org. Quarterly releases from DepMap are continually updated and integrated into DDH. At the time of writing this manuscript, the 20Q1 release was used, which contains gene essentiality scores for 17426 genes across 739 cell lines. Current data release can be found at datadrivenhypothesis.org

### Data cleaning

A permutation test typically involves permuting one or more variables in a data set before performing the test, in order to break any existing relationships and simulate a null hypothesis of random sampling. In this case, we broke the relationship between gene-gene pairs and the correlation values. We then generated a distribution of null statistics (fake mean correlations), along with standard deviations of these sampled data. This strategy identified at which correlation value to draw a threshold of a “significant correlation” for these analyses.

### Pubmed

PubMed contains 30 million publication records containing article metadata and abstracts. We began by first parsing each of the 30 million abstracts for all gene names and gene aliases in an abstract. Next we were able to generate article level summaries for each gene in each article, and then counted gene-gene co-occurrence in an abstract. Measuring all human gene-gene co-occurrences across all PubMed records allowed us to identify the frequency with which a gene was published with another gene. Unfortunately, gene nomenclature is a mess, with official gene names changing over time, and several genes sharing common aliases. As a keystone example, the protein p38 has been used as a name to describe the genes MAPK14 (NCBI gene id 1432), AHSA1 (NCBI gene id 10598), and AIMP2 (NCBI gene id 7965). The protein p38 associated with MAPK14 is among the most studied and published on genes in the human genome, whereas AHSA1 and AIMP2 are among the least studied. This discrepancy and ambiguity in identifying genes in abstracts necessitated an alternative approach. Therefore, we leveraged a PubMed resource from the National Library of Medicine (NLM) called gene2pubmed that links unique NCBI gene ids to a PubMed id. This resource is part of the NLM’s Indexing Initiative (IND), which is working to automate manual indexing practices. Using gene2pubmed, we again measured all human gene-gene co-occurrences across all PubMed records. While this method overcomes the challenges associated with programmatically assigning gene names by providing gene ids, it is still limited in its ability to curate genes associated with scientific papers by the same challenges.

## Data Availability

The Broad Institute has information on their website and the website dedicated to the Dependency Map project about how the raw data were generated, and provide a list of references. DDH is the resource leveraging these data for identifying novel functions for human genes developed by the Hirschey Lab. Code is available on the Hirschey Lab github account (www.github.com/matthewhirschey/ddh), including links to download the raw data, and run the analyses in R from scratch.

## Usage

https://www.datadrivenhypothesis.org

---

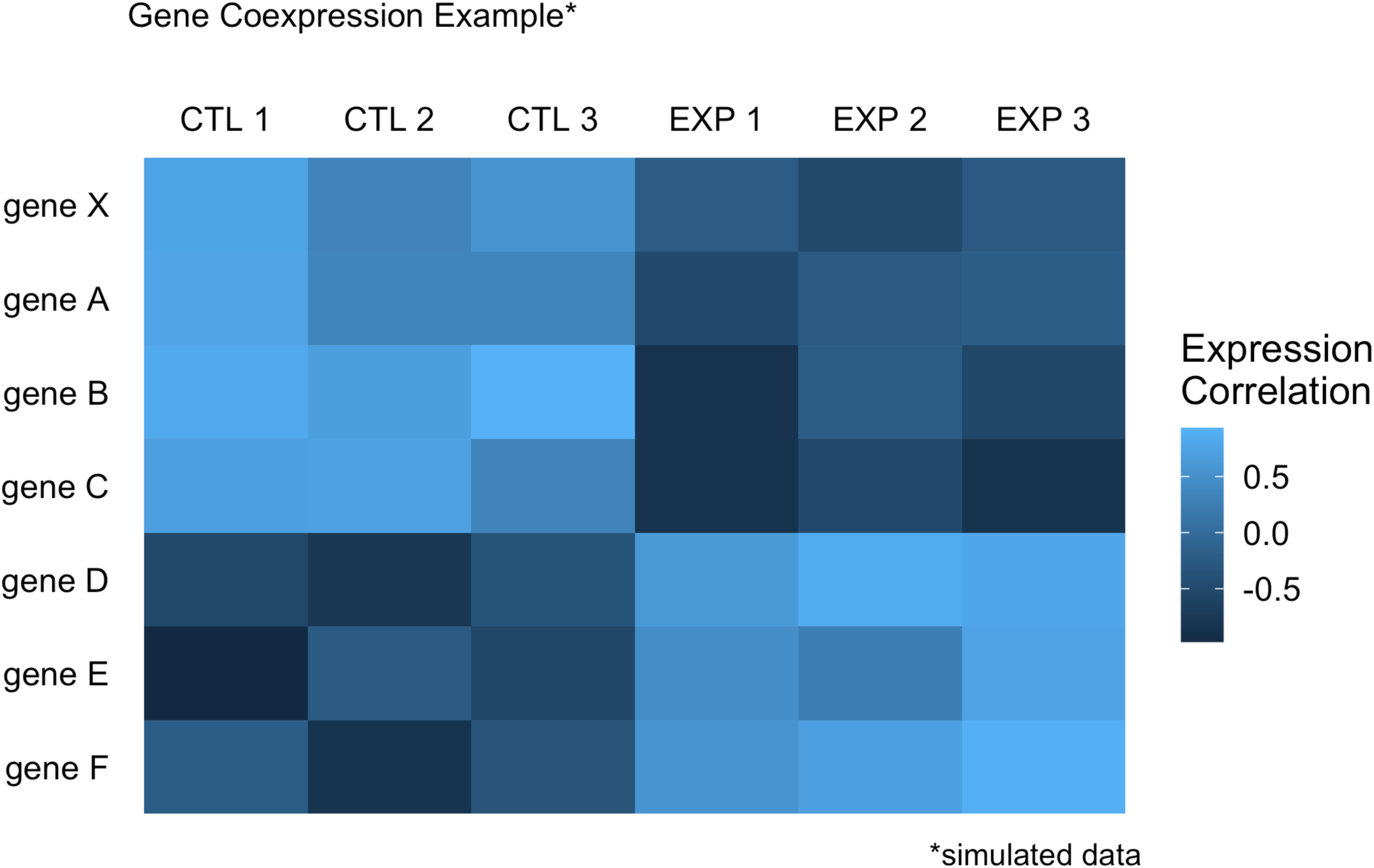

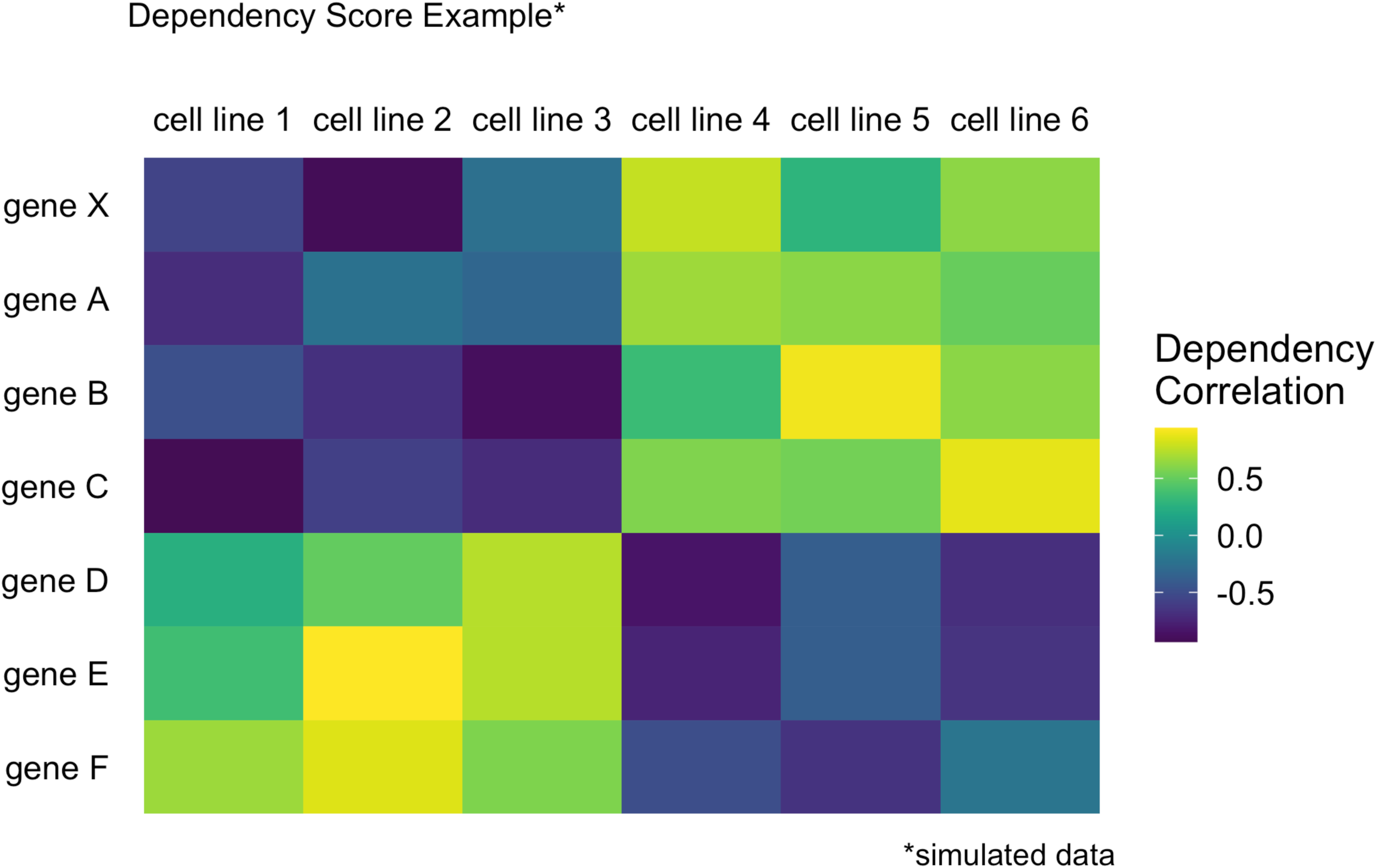

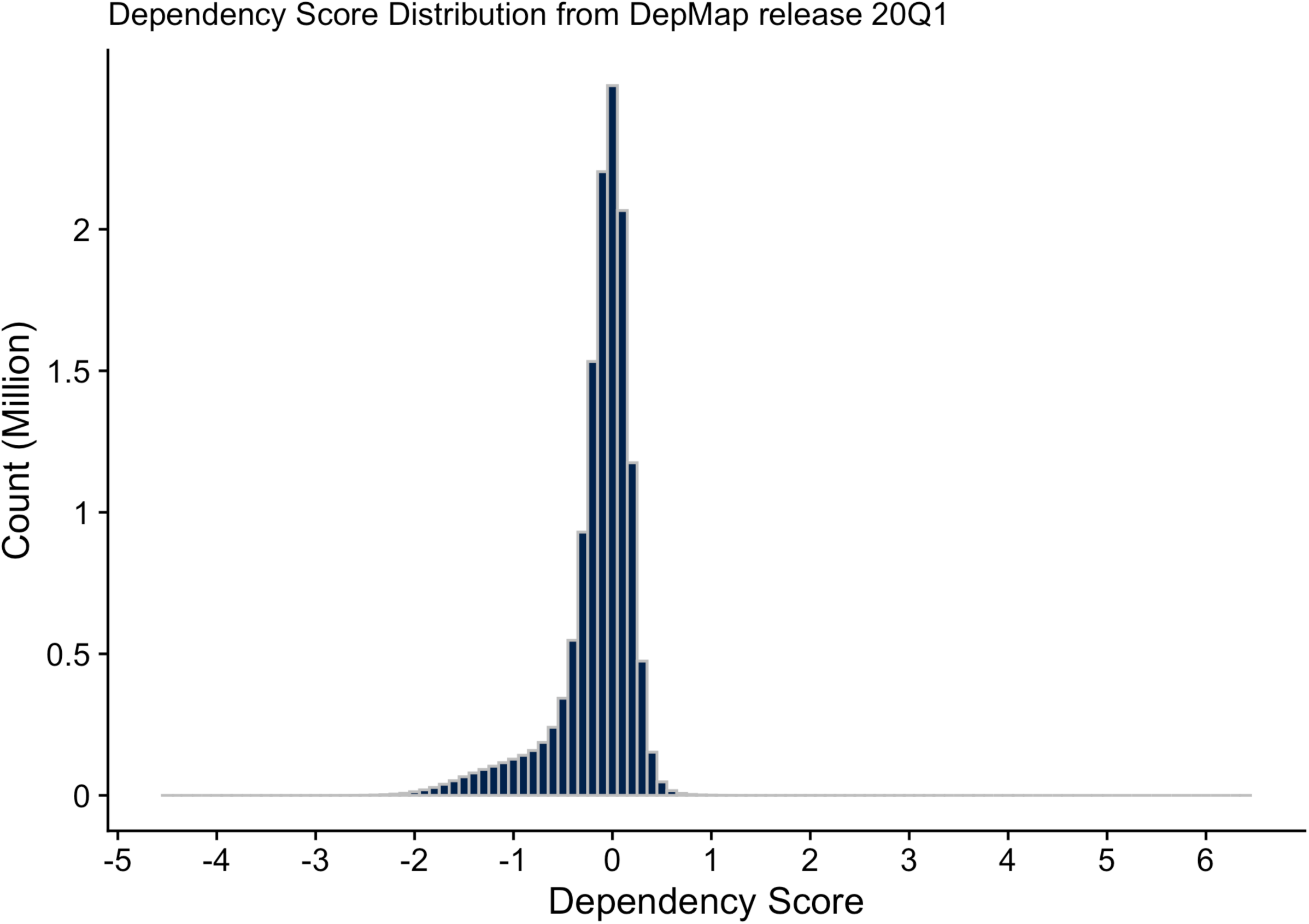

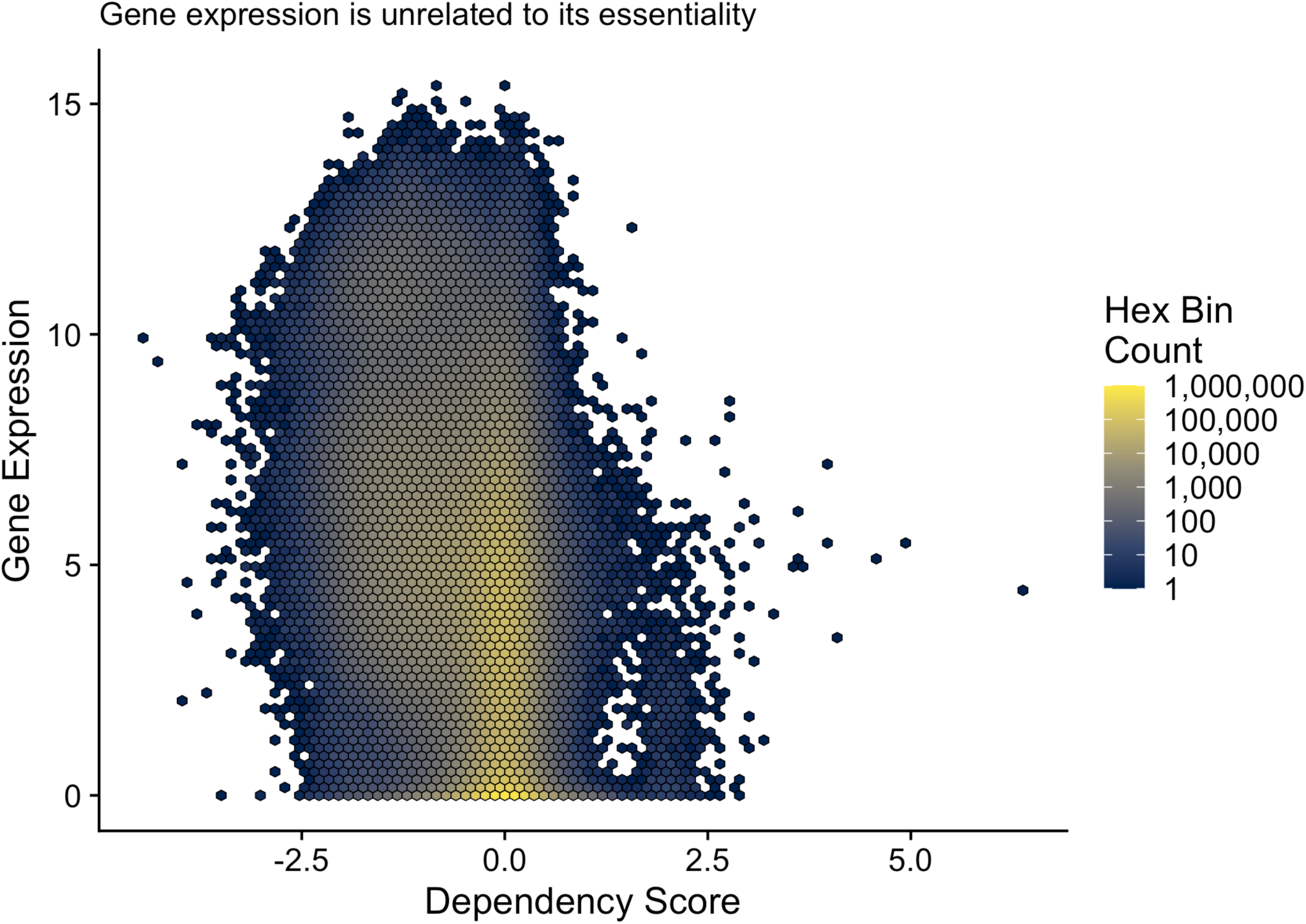

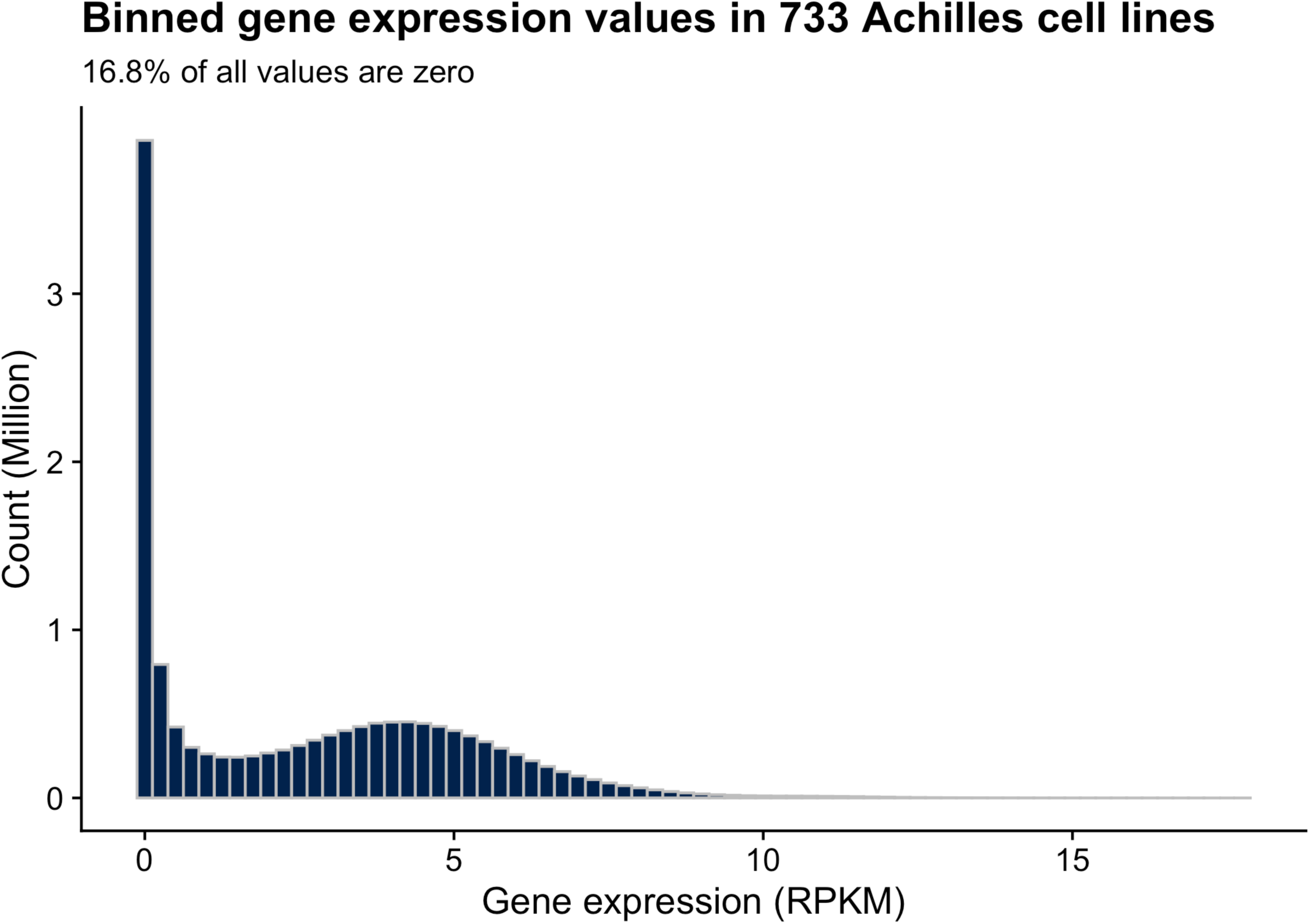

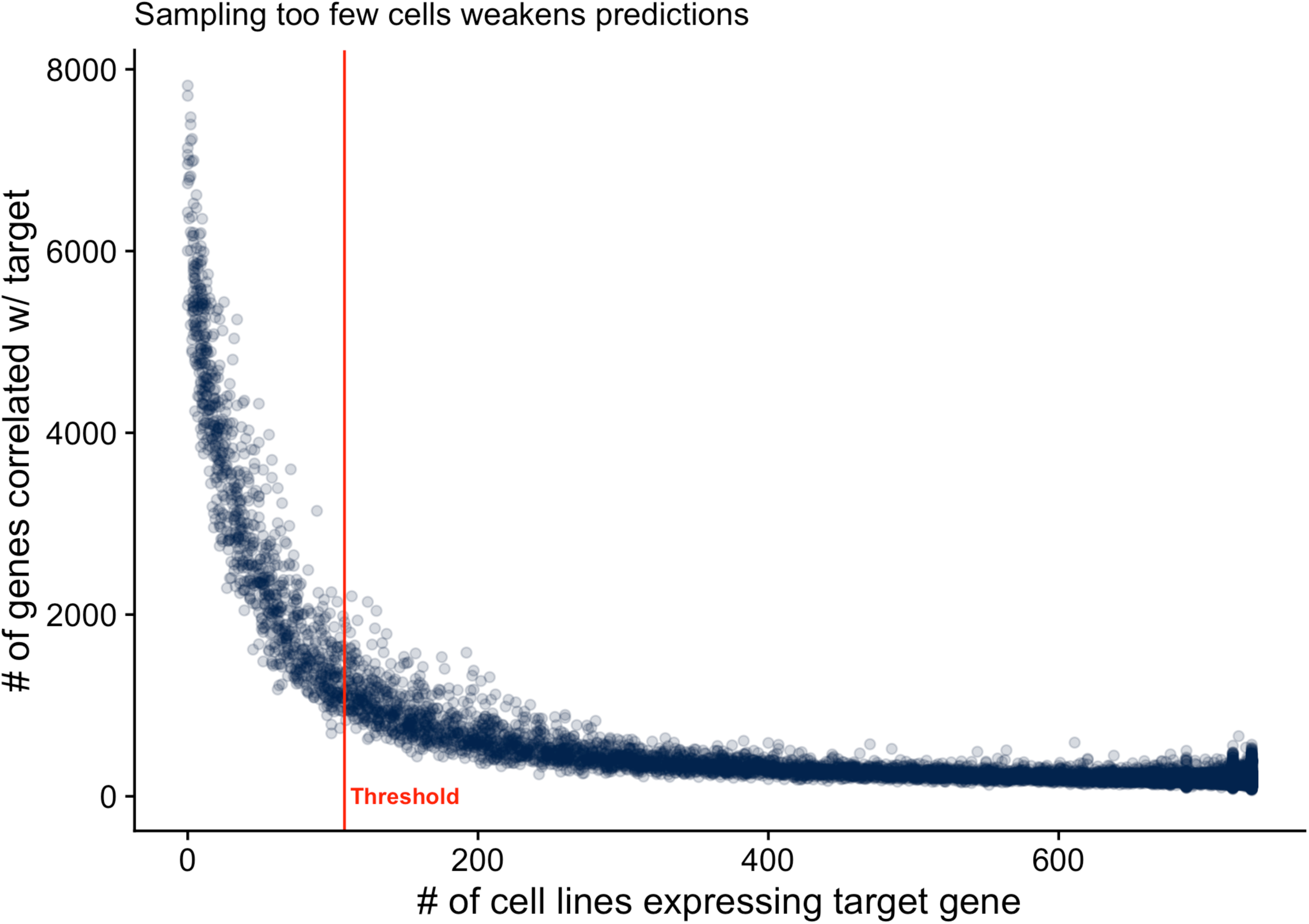

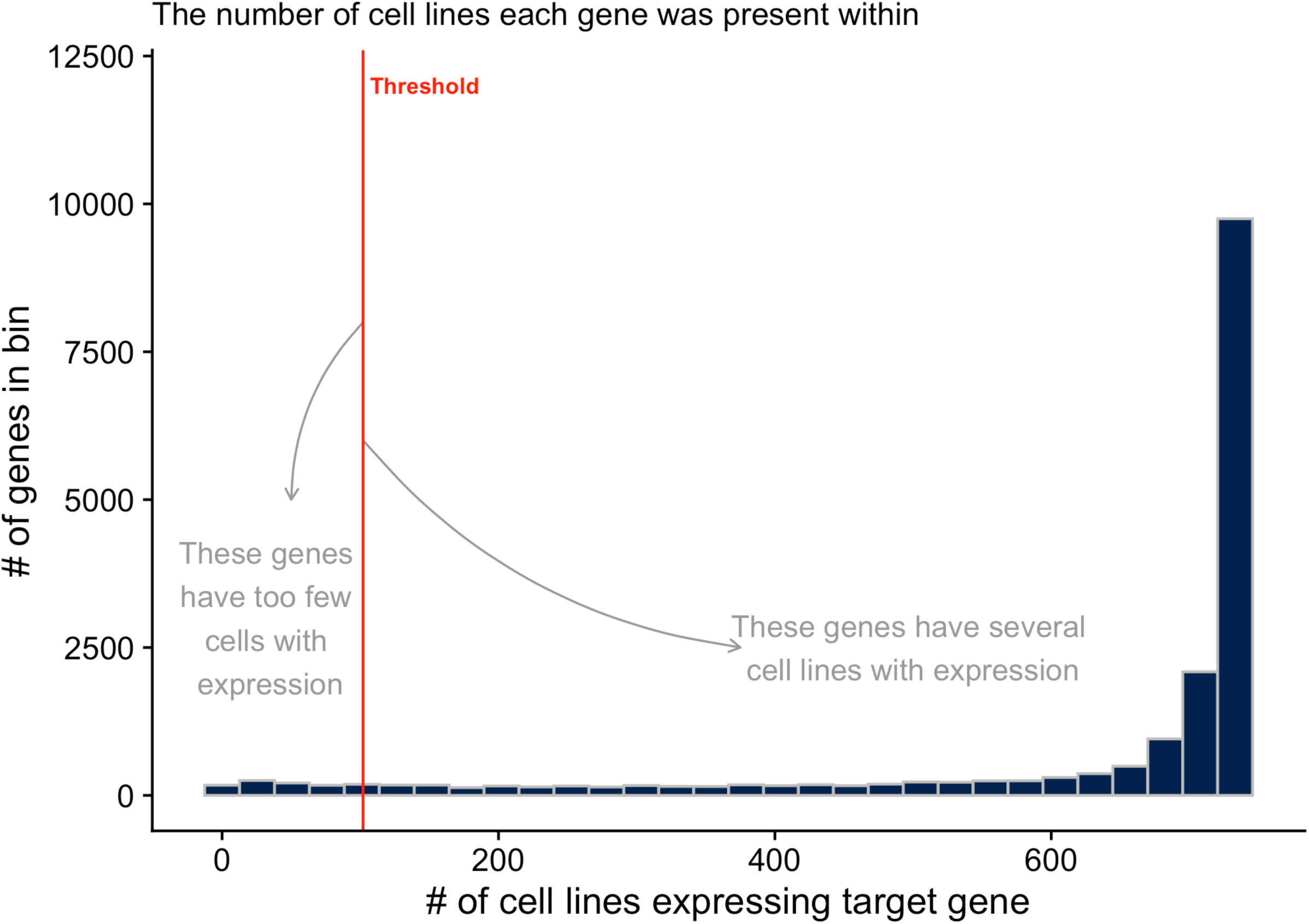

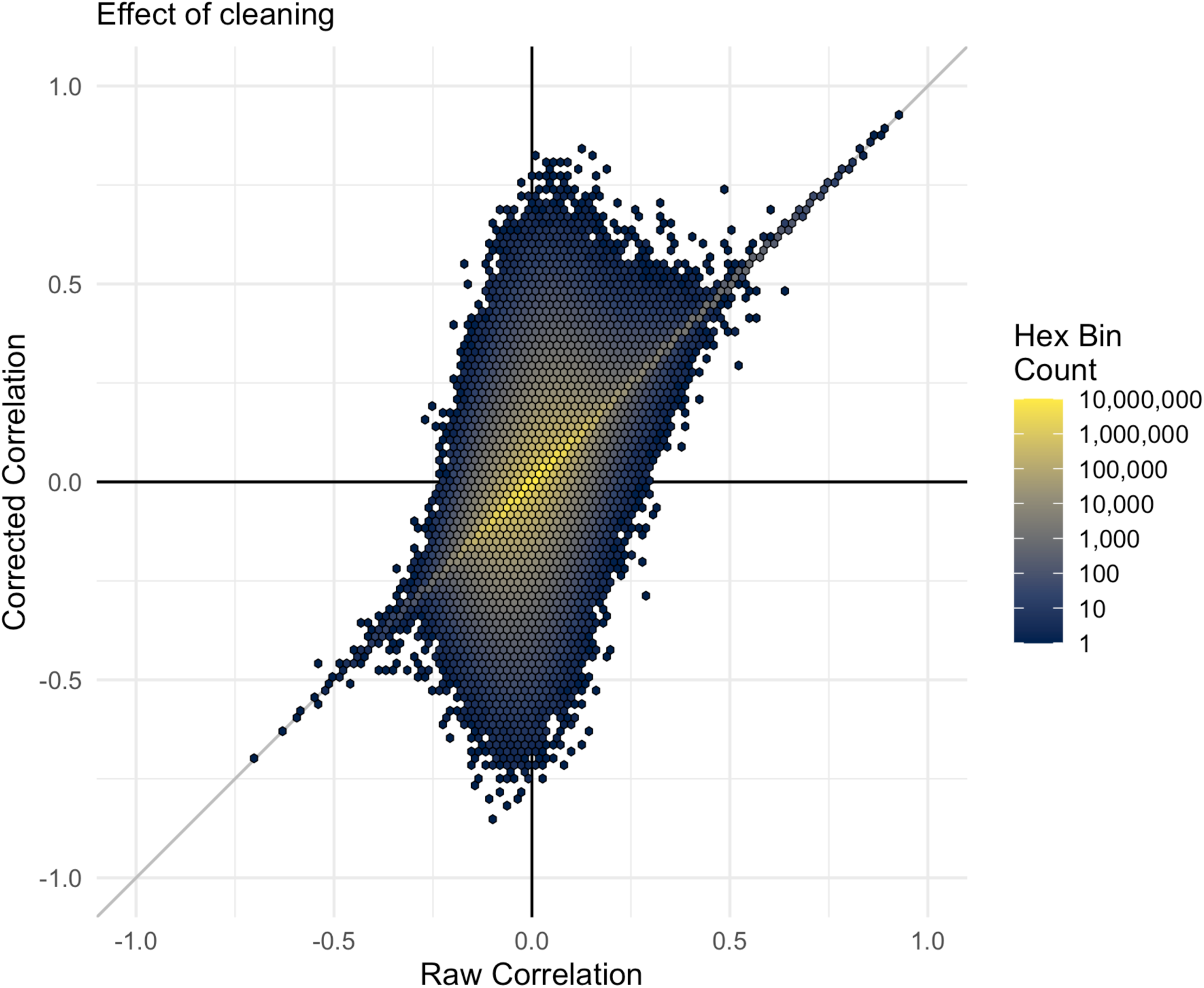

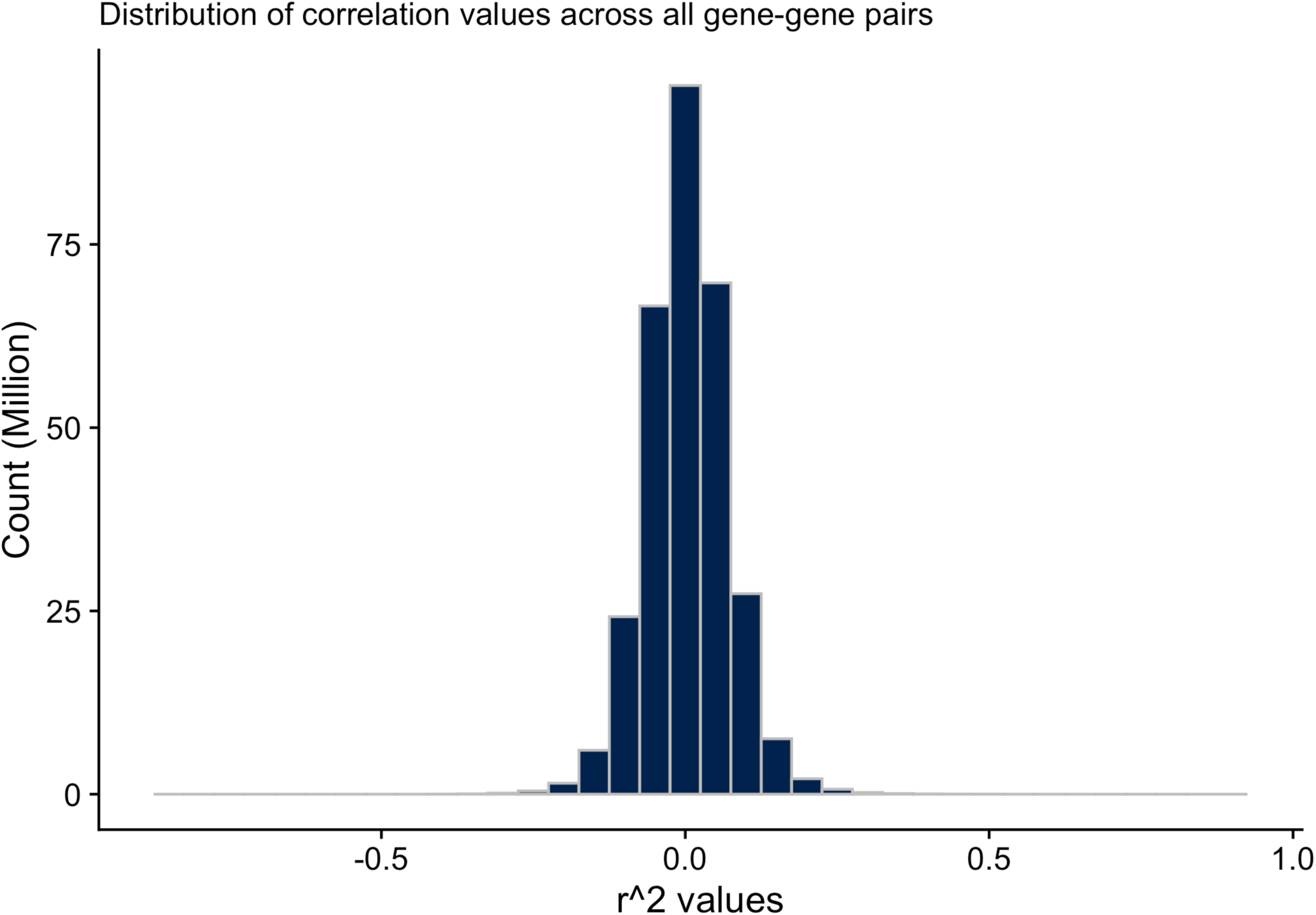

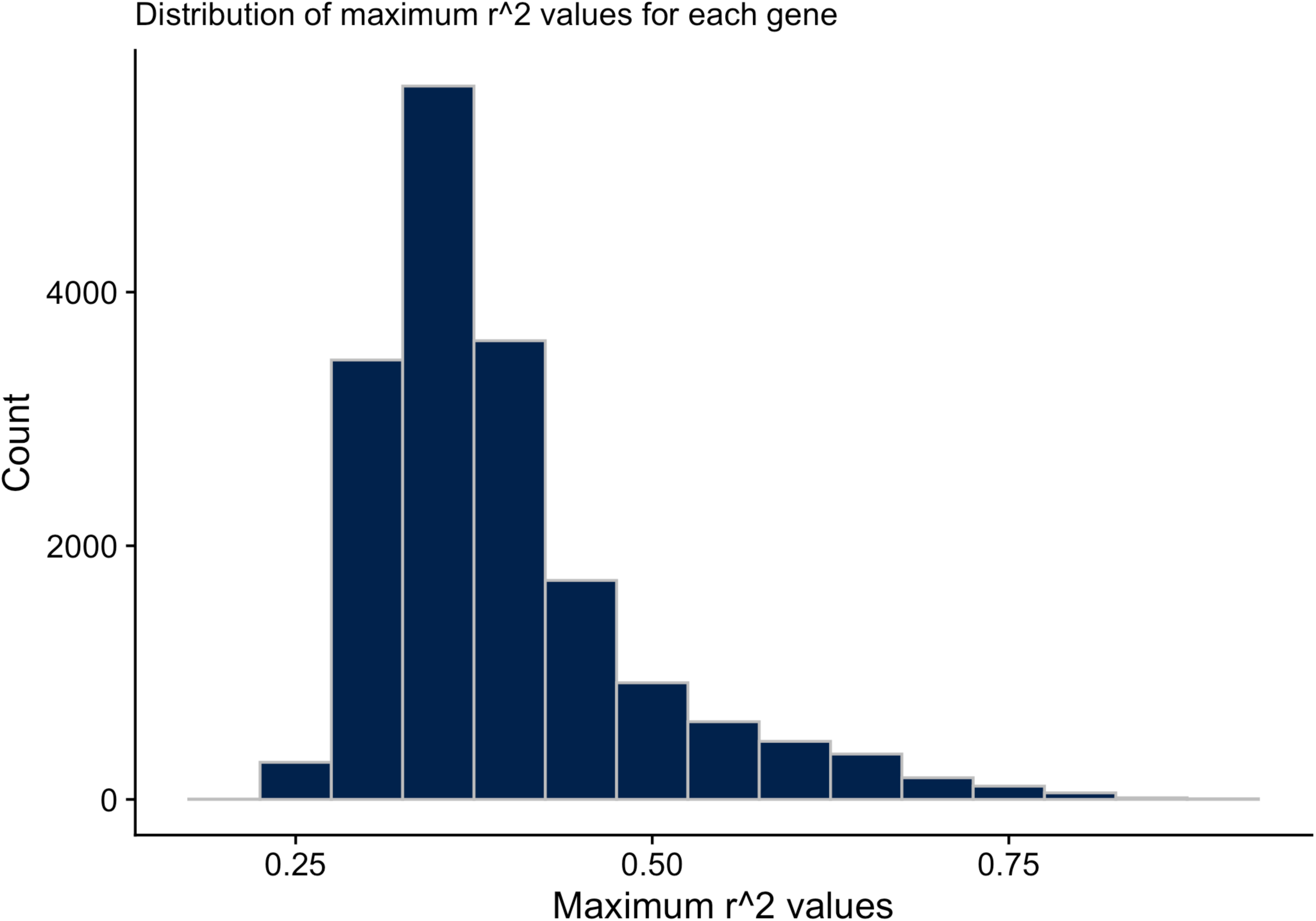

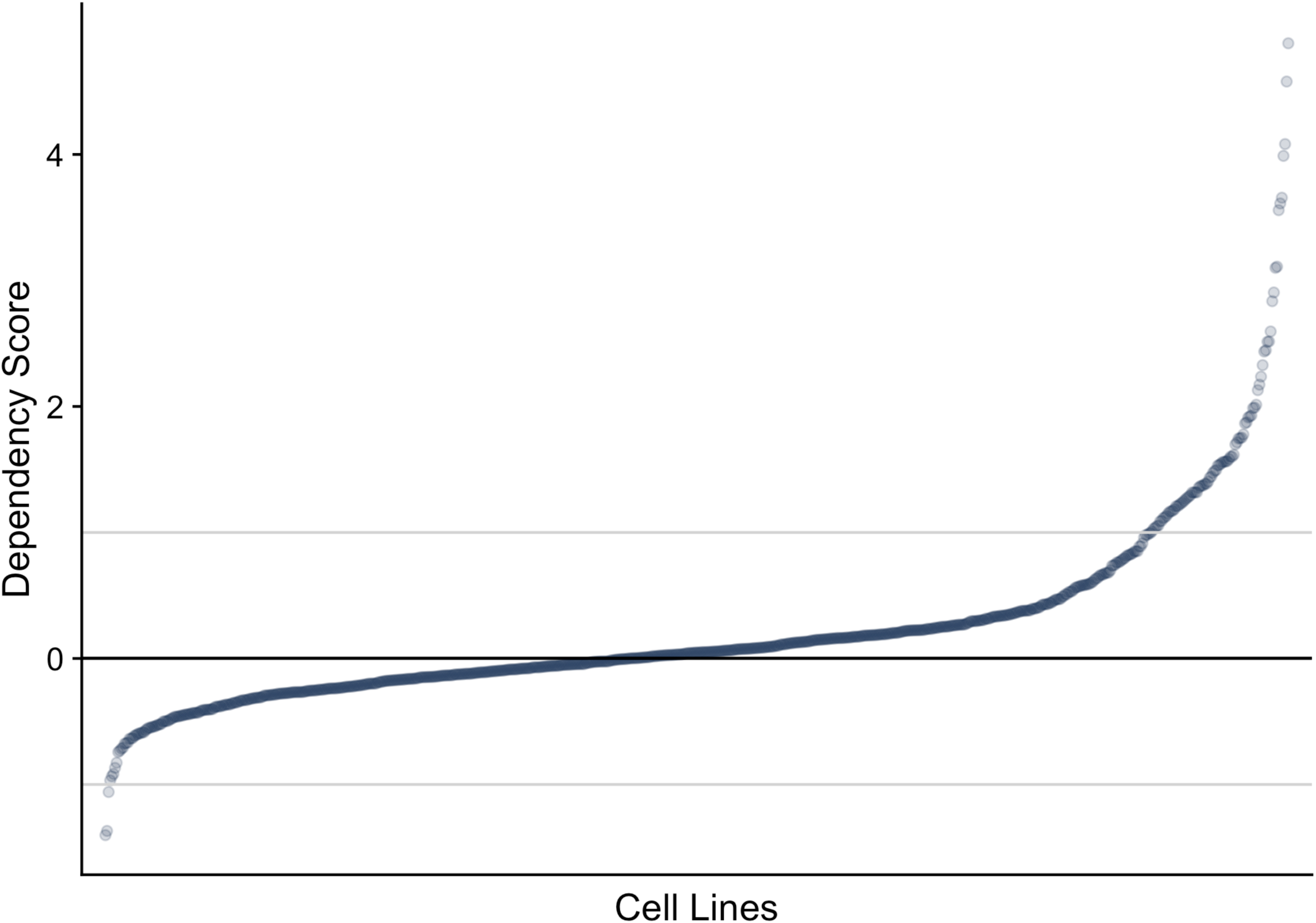

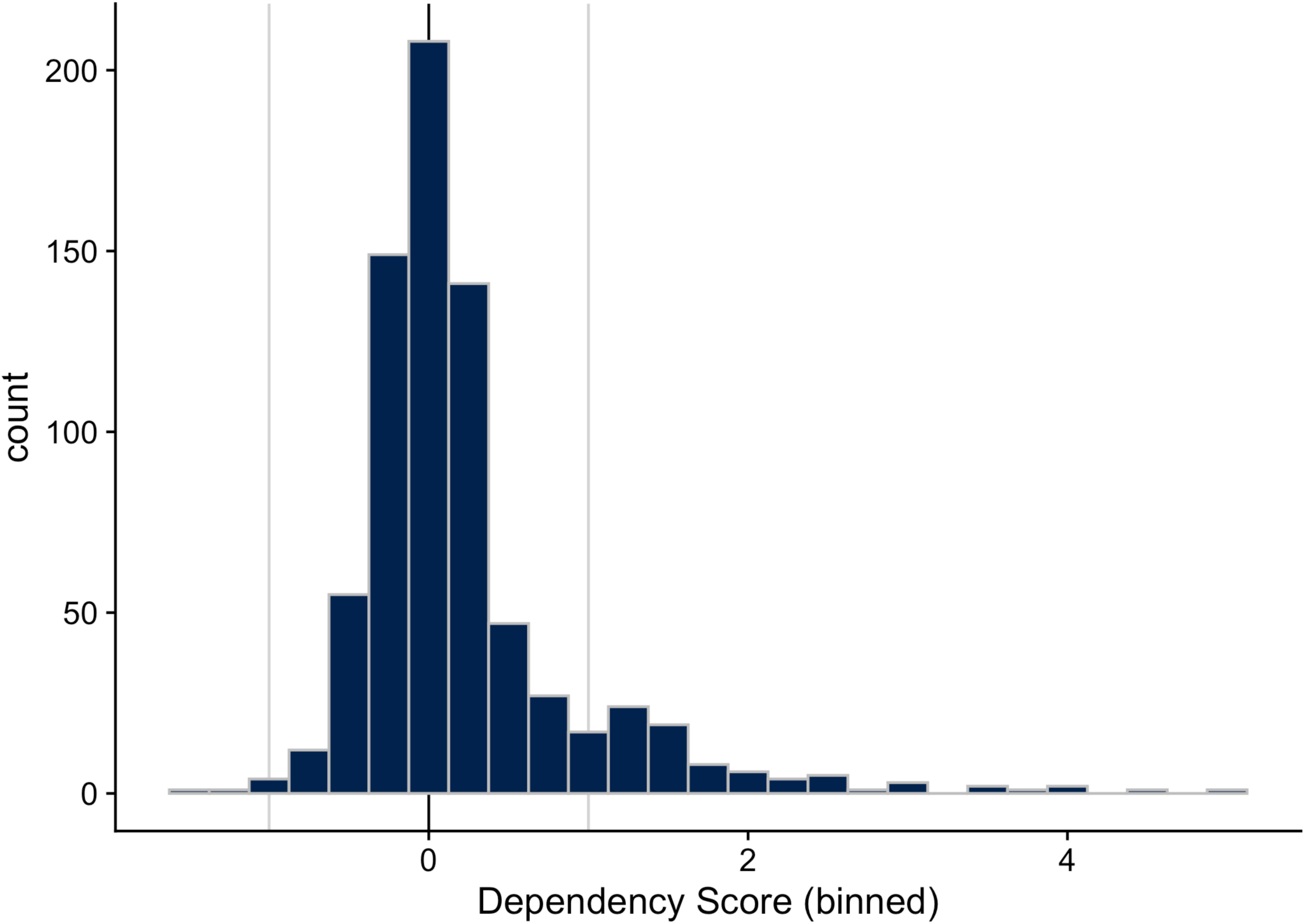

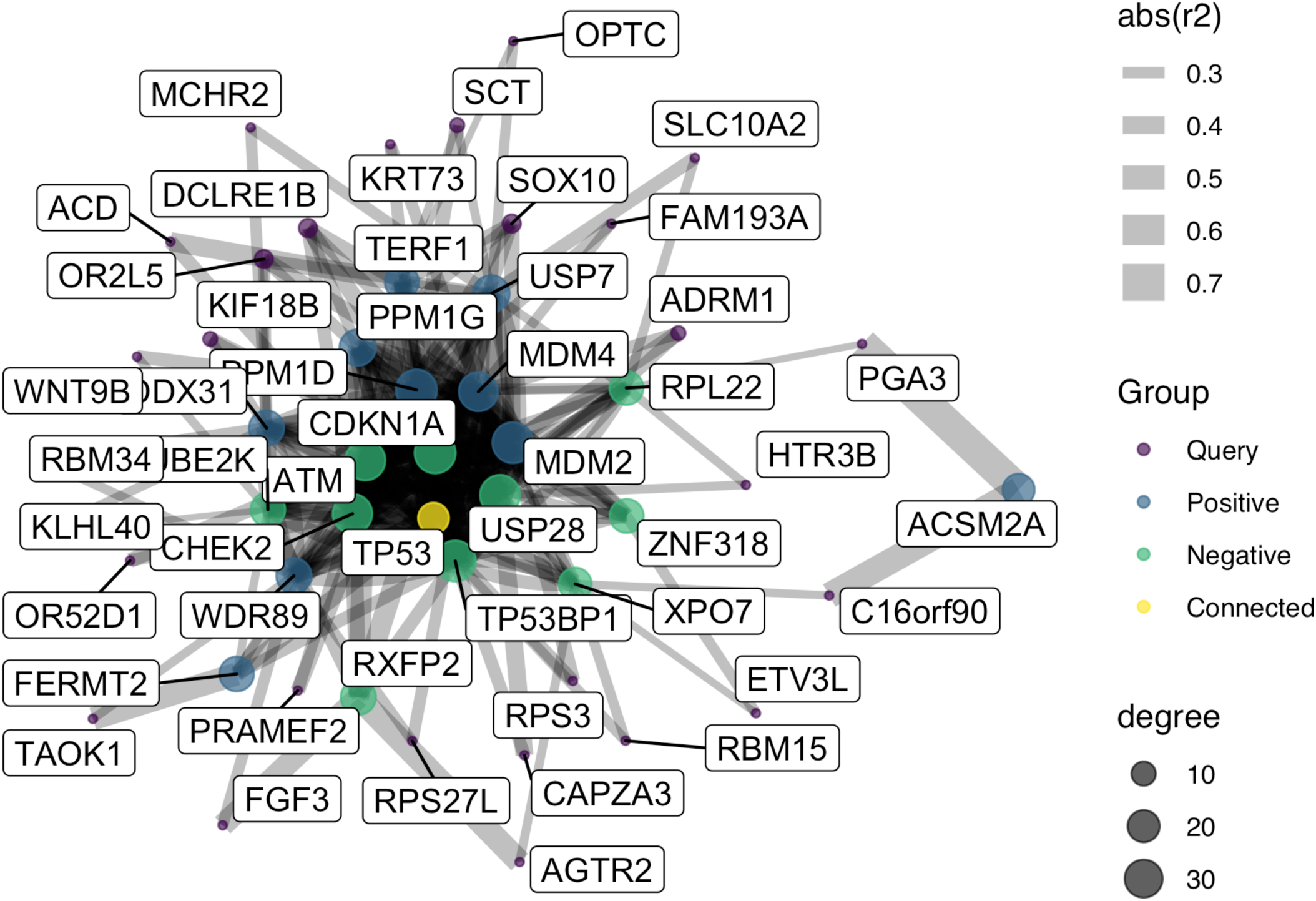

